# Transposon-triggered epigenetic chromatin dynamics modulate EFR-related pathogen response

**DOI:** 10.1101/2023.10.06.561201

**Authors:** Regina Mencia, Agustín L. Arce, Candela Houriet, Wenfei Xian, Adrián Contreras, Gautam Shirsekar, Detlef Weigel, Pablo A. Manavella

## Abstract

Infectious diseases drive the evolution of wild plants and impact yield in crop plants. Like animals, plants can sense biotic threats via conserved pathogen-associated patterns (PAMPs). Since an overly robust immune response can harm plants, understanding the mechanisms for tuning defense responses to the appropriate level is vital as we endeavor to develop pathogen-resistant crops. In this paper, we studied the Arabidopsis pattern recognition receptor (PRR) EFR, which senses bacterial EF-Tu. An inverted-repeat transposon (*Ea-IR*) between *EFR* and the neighboring *XI-k* locus controls local chromatin organization, promoting the formation of a repressive chromatin loop. Upon pathogen infection, the chromatin landscape around *EFR* and *Xl-k* dynamically changes to allow for increased *EFR* transcription. Chromatin opening facilitates the passage of RNA polymerase II across the neighboring *XI-k* gene termination site, leading to a longer *XI-k* transcript that includes *Ea-IR* sequences. Dicer-like (DCL) enzymes process the longer Xl-k transcript into small RNAs (sRNAs), which reset chromatin topology to a repressive state, attenuating, in turn, the immune response, reminiscent of attenuation of receptor signaling in other systems. From an evolutionary point of view, we found that natural Arabidopsis accessions missing *Ea-IR* have a constitutive "*EFR*-open" chromatin configuration that correlates with higher basal EFR levels and higher background resistance to pathogens. Collectively, our study offers evidence for a scenario in which a transposon, chromatin organization, and gene expression interact to fine-tune immune responses, both during the course of infection and in the course of evolution. Similar gene-associated IRs in crops could provide valuable non-coding targets for genome editing or assisted plant breeding programs.

## Introduction

Like vertebrates, plants have evolved complex mechanisms in which two arms of the immune system interact to efficiently combat pathogens while minimizing collateral damage to the host ^1^. The first layer, akin to animal innate immunity, employs pattern-recognition receptors (PRRs) situated on cell surfaces to sense conserved pathogen (or microbe)-associated molecular patterns (PAMPs/MAMPs), initiating PAMP-triggered immunity (PTI) ^2,3^. The second layer involves intracellular receptors that recognize pathogen-released effectors, activating effector-triggered immunity (ETI), resulting in a hypersensitive response and programmed cell death ^1,2^. Nucleotide-binding site leucine-rich repeat proteins (NLR or NBS-LRR), many of which were identified genetically as encoded by major Resistance (R) genes, act as principal ETI immune receptors^4,5^. PTI and ETI, often synergize to optimize plant immunity ^6-10^.

During PTI, activation of PRRs initiates a series of signal transduction events leading to changes in gene expression and increased basal disease resistance against pathogens that are not adapted to the host ^3,11,12^. Most plant PRRs are receptor kinases (RKs) or receptor-like proteins (RLPs) with extracellular ligand-binding, transmembrane, and intracellular kinase domains ^11^. The Brassicaceae-specific PRR ELONGATION FACTOR-TU RECEPTOR (EFR) recognizes bacterial EF-Tu ^13-18^. Increased pathogen resistance often correlates with elevated *EFR* expression ^19-23^.

Many organisms, from prokaryotic and eukaryotic microbes to invertebrates and vertebrates, can harm or kill plants in nature. However, while it is imperative for plants to promptly activate defense responses when attacked, these must be tuned to the size of the threat to avoid collateral damage in the form of reduced growth and seed set ^24,25^. Hence, highly effective immune alleles have an advantage in populations under high pathogen pressure but decline in non-threatening environments. Indeed, both PRR and NLR genes are often polymorphic in natural populations of flowering plants ^26-29^. This is particularly true for NLRs, because many of the pathogen effectors recognized by NLRs are not essential for pathogen survival ^30^. Conversely, the homeostatic function and high conservation of most PAMPs cause PRR receptors to be less genetically variable ^12^. Nevertheless, apart from the absence/presence variation of specific genes or alleles, variation in transcript or protein levels can influence PRR activity and alter immune responses in natural populations ^31^. Because *PRR* RNA levels are known to be positively regulated ETI, this indicates that there are likely opportunities for fine-tuning *PRR* expression levels before and during pathogen attack ^32^.

Differences in gene expression can be caused *in trans* by accession-specific transcriptional regulators or by *cis*-regulatory sequence variants. Recently, we found that the insertion of TE-derived inverted repeats (IRs) near genes changes the local chromatin organization, with profound impacts on the activity of neighboring loci ^33,34^. Mechanistically, transcription of these IRs leads to the synthesis of 24-nt small RNAs and the methylation of the IRs. This provides an anchor point for the formation of short-range chromatin loops that influence the expression of neighboring genes ^35^. Many IRs vary between *A. thaliana* accessions (with many accessions missing the entire IRs), causing differences in short-range chromatin interactions and gene expression ^34^. Such variation in the gene-hosted IR content is significant because it can be associated with changes in phenotypic traits ^34^. This points to IRs as powerful evolutionary agents that can epigenetically contribute to plant adaptation. In this sense, while TEs are commonly silenced through the RNA-dependent DNA Methylation (RdDM) pathway that relies on the plant-specific RNA polymerases IV and V (Pol IV and Pol V) and the RNA-dependent RNA Polymerase 2 (RDR2) ^36,37^, IRs can trigger DNA methylation by a non-canonical pathway as Pol II transcripts of IRs can fold into dsRNAs triggering sRNA production without the need of RDR2 ^38^. This turns IRs into autonomous elements that, when inserted near a gene, can be transcribed, triggering local RdDM, chromatin looping, and changes in gene expression ^34^.

Here we show that an IR that separates *EFR* from the myosin XI-k gene (*Xl-k*), which associated with root organogenesis, ER dynamics, and fungal penetration ^39-41^, controls *EFR* expression by shaping local chromatin topology. Mechanistically, the methylation of the IR induces the formation of a chromatin loop that represses *EFR* expression. Upon pathogen infection, demethylation of the TE causes the opening of the loop and the activation of *EFR* transcription. Methylation of the IR is maintained by canonical Pol IV-dependent RNA-directed DNA methylation (RdDM), initiated by the processing of a long, IR-containing *XI-k* transcript. This long transcript results from a Pol II read-through event traversing the termination site when the loop is in the open conformation, as is the case during pathogen infection. The IR is found in only some *A. thaliana* accessions. Accessions without the IR present a distinctive loci topology, higher basal levels of EFR, and enhanced resistance to pathogens. Due to the central role of EFR in plant innate immunity, this naturally occurring epigenetic variation gain relevance for environmental adaptability and evolution.

## Results

### An IR between *EFR* and *XI-k* provokes chromatin looping and *EFR* expression

Because of the recently discovered link between structural variants involving IRs near genes and potentially adaptive phenotypic variation in *A. thaliana* natural accessions ^34^, we searched for IR proximal genes with roles in pathogen response. We focused on IRs within 500 bp of protein-coding genes, as these IRs tend to have the most profound impact on chromatin organization and gene expression ^34^. Among the 261 genes located ≤500 bp from an IR element, the overlap between Gene Ontology (GO) categories "response to stress" and "response to biotic stimulus" included 24 genes (Figure S1A), including the well-known PRR gene *EFR* (AT5G20480) and the adjacent *XI-k* (AT5G20490) gene, which also has been linked to pathogen-relevant biological processes. A 442 bp long IR occupies almost the entire 615 bp intergenic region separating these two pathogen-related genes (Figure 1A). This IR, named *EFR-associated IR* (*Ea-IR*) for simplicity, corresponds to an element of the ATMU3N1 family of DNA/MuDR transposons and is annotated as a miniature inverted-repeat transposable element (MITE) in the P-MITE database ^42^.

**Figure 1.**
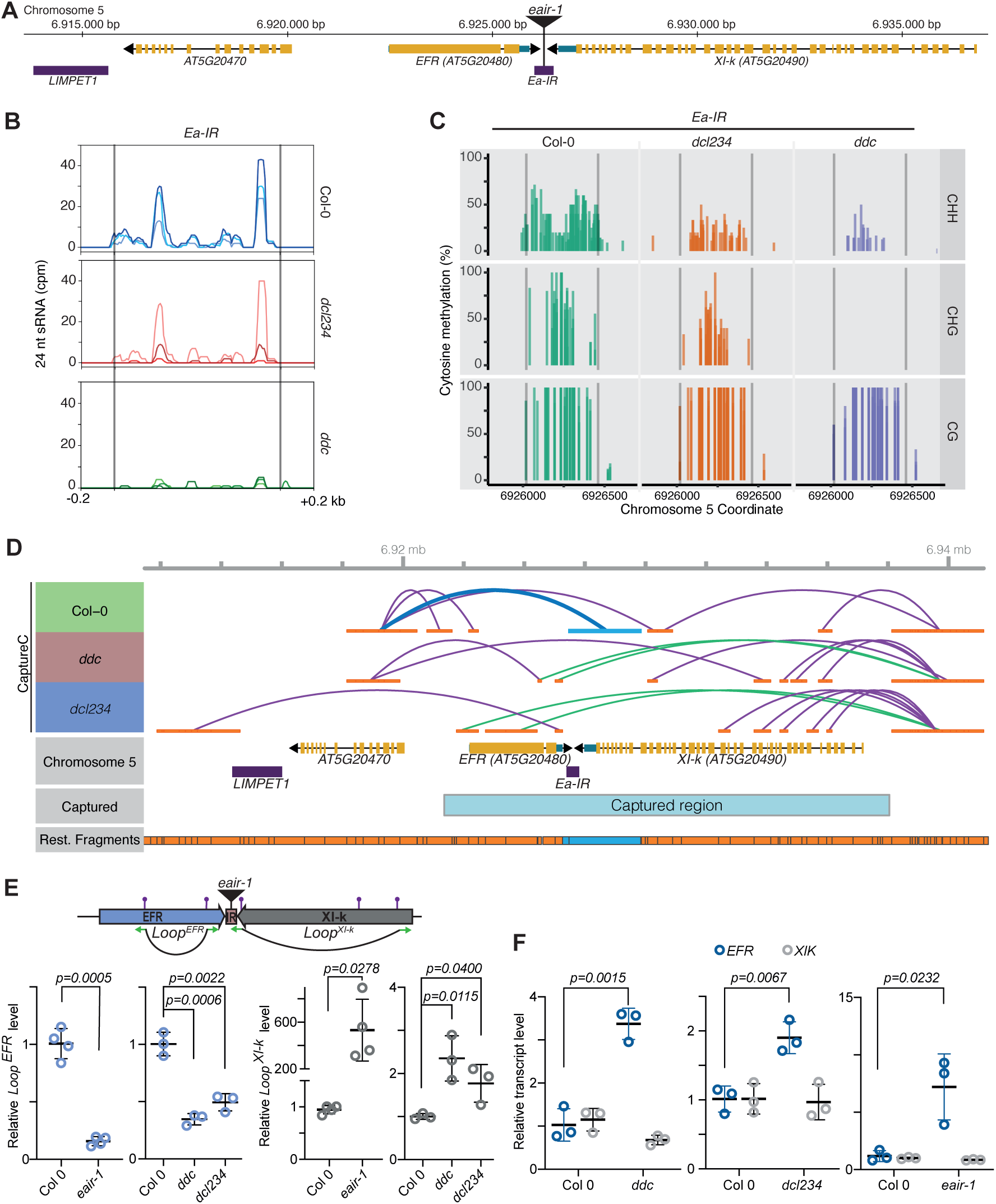
*Ea*-*IR* methylation affects *EFR* expression. (A) Diagram of the *EFR*/*Xl-k* region on chromosome 5 (yellow bars exons, blue bars UTRs). The *LIMPET1* transposon and the transposon-derived *Ea-IR* inverted repeat are indicated by purple bars. The T-DNA insertion disrupting *Ea-IR* in the *eair-1* mutant is also indicated. Numbers = chromosome 5 coordinates. (B) 24 nt siRNAs mapping *Ea-IR* in Col-0 wild-type, *dcl234*, and *ddc* plants. For each genotype, the results from three replicates are shown in different shading. (C) Cytosine methylation in the CHH, CHG and CG contexts at *Ea-IR* (delimited by vertical gray lines) in Col-0 wild-type, *dcl234*, and *ddc* plants. Three individual BS-seq replicates are plotted for each genotype with different shades. (D) Chromatin contacts from CaptureC in the *EFR/Xl-k* region in Col-0 wild-type, *ddc,* and *dcl234* plants. *Loop^EFR^*, which includes *Ea-IR*, is marked in cyan, *Loop^Xl-k^* in green, and all other interactions in purple. The region targeted by CaptureC is indicated, with all potential restriction fragments below. Fragments diagnostic for the *Ea-IR* region are indicated in cyan. (E) On top, purple pins indicate XbaI sites used for 3C experiments. Given the resolution of 3C experiments, the anchor points for the two loops, which are indicated as well, can be located anywhere within each of the ligated restriction fragments (green arrow). On the bottom, quantification of *Loop^EFR^* (blue rings) and *Loop^Xl-k^* (gray rings) by 3C-qPCR in Col-0 wild-type, *eair-1*, *ddc*, and *dcl234* plants. (F) Expression of *EFR* (blue rings) and *Xl-k* (gray rings) as measured by qRT-PCR and normalized against *ACT2*. In E and F, values of n = 4 biologically independent mutant samples are expressed relative to the average values of Col-0 wild-type plants. Data are presented as mean values ± s.d. *P* values were calculated with a two-tailed unpaired t-test with Welch’s correction.

IRs can produce 24 nt long siRNAs that trigger DNA methylation at the IR that spawns them, leading to changes in chromatin organization and expression of adjacent genes ^33,34^. Abundant 24 nt siRNAs were detected at *Ea-IR* (Figure 1B), as was methylation in all cytosine contexts (Figure 1C and S1C). CHH and CHG methylation and siRNAs are significantly reduced in *drm1 drm2 cmt3* (*ddc*) and *dcl2 dcl3 dcl4* (*dcl234*) mutants, indicative of involvement of the RdDM pathway (Figure 1B and 1C). Abundant 24 nt siRNAs and DNA methylation were also found at a 2,060 bp *LIMPET1* DNA/MuDR transposon located 6,620 bp upstream of *EFR* and 480 bp downstream of AT5G20470, a headless derivative of *XI-k* (Figure 1A, S1C-E). Given the non-IR nature of this *LIMPET* transposon and its proximity to *Ea-IR*, it served as a valuable control to discern specific IR-triggered molecular effects in subsequent analyses.

To determine whether *Ea-IR* affected local chromatin organization, we analyzed published Capture-C data ^34^, which revealed a short-range chromatin loop that included the entire *EFR* coding region (Figure 1D, dark blue interactions). This loop, termed *Loop^EFR^*, appeared to be disrupted in *ddc* and *dcl234* mutants, suggesting that its formation relies on *Ea-IR* methylation (Figure 1D). In contrast, a second loop encompassing the *XI-k* gene (*Loop^XI-k^*) was only detected in the RdDM-deficient mutants (Figure 1C, green interactions). These findings are in agreement with a role of siRNA-induced DNA methylation in forming and stabilizing specific short-range chromatin loops.

To study *Loop^EFR^* and *Loop^XI-k^* and their dependence on *Ea-IR* in more detail, we conducted Chromatin Conformation Capture followed by qPCR (3C-qPCR) in Col-0 wild-type plants, *ddc* and *dcl234*, and a T-DNA insertion line (WiscDsLox440E09), hereafter referred to as *eair-1*, which contains a large insertion in the middle of *Ea-IR* (Figure 1A and S1B). These experiments demonstrated that the chromatin loop encompassing the *EFR* gene in wild-type plants was dependent on an intact and methylated *Ea-IR* (Figure 1E). Collectively, our data reinforce the idea of *Loop^EFR^* formation triggered by *Ea-IR* in combination with the RdDM pathway (Figure 1E). That the formation of *Loop^XI-k^* was enhanced in *eair-1* and RdDM-deficient mutants, which lack the *Loop^EFR^*, implies competition between the two loops, with *Loop^EFR^* being dominant (Figure 1E). We repeated the 3C-qPCR analyses with different sets of restriction enzymes to more precisely delineate the interacting regions of *Loop^EFR^* and *Loop^XI-k^*. These experiments confirmed that *Loop^XI-k^* only forms in the absence of *Loop^EFR^* (Figure S2).

Finally, to assess whether changes in chromatin organization caused by physical disruption of *Ea-IR* in *eair-1* mutants or by demethylation of *Ea-IR* in RdDM deficient mutants affected the expression of nearby genes, we measured *EFR* and *XI-k* mRNA accumulation. Only *EFR*, but not *XI-k*, was more highly expressed in the mutants tested (Figure 1F), defining *Loop^EFR^* as a repressive loop. That *XI-k* expression was unaltered indicated that *Loop^EFR^*effects on *EFR* are specific, as *Ea-IR* is similarly close to both *EFR* and *Xl-k*.

### Chromatin loops in the *EFR/XI-k* region are sensitive to activation of immunity

Upon pathogen infection, many regions in the *A. thaliana* genome experience a reduction in DNA methylation levels, which is often coupled to increased production of 21-nt siRNAs ^43,44^. This phenomenon correlates with heightened pathogen resistance in plants lacking core elements of the RdDM pathway ^45^, and conversely, reduced resistance in mutants lacking the important cytosine demethylase ROS1 ^46^. In accordance, we observed that DNA methylation at the *Ea-IR* is markedly reduced upon *Pseudomonas syringae* pv. tomato strain DC3000 *hrcC* (*Pst* DC3000 *hrcC*), a type III secretion system-deficient strain that facilitates the detection of PTI-related phenotypes independently of the activation of ETI ^47^. (Figure 2A). We posited that this pathogen-induced alteration of the epigenetic landscape could potentially modify chromatin organization and expression of the *EFR* locus through effects on *Ea-IR*. To investigate this hypothesis, we infected Col-0 plants using *Pst* DC3000 *hrcC*, Infection with *Pst* DC3000 *hrcC* induced the expression of *EFR* and the defense-associated marker *PATHOGENESIS-RELATED GENE 1* (*PR1*), but not of *XI-k* (Figure 2B).

**Figure 2.**
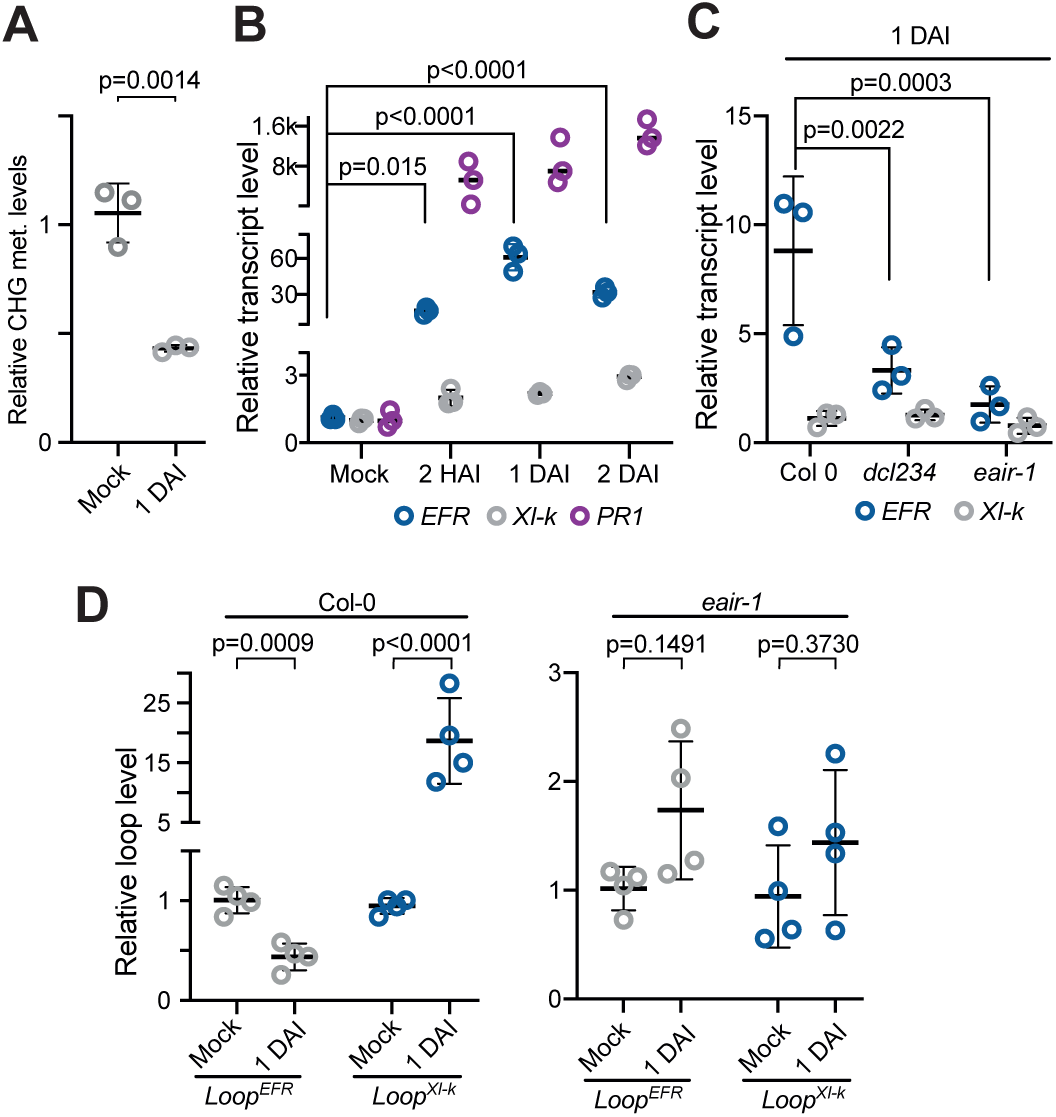
Pathogen infection alters chromatin architecture and gene expression at the *EFR/XI-k* locus. (A) MspI Chop-qPCR analysis of *Ea-IR* DNA methylation in three biological replicates of Col-0 wild-type plants mock-treated or after one-day infection with *Pst* DC3000 *hrcC* (1 DAI). Digestion efficiency was quantified by qPCR with primers spanning restriction sites in the sRNAs mapping regions and normalized to an undigested region. (B and C) Relative expression of *EFR*, *Xl-k*, and *PR1* (only in A) in Col-0 wild-type (B), and *dcl234* and *ddc* (C) plants infected with *Pst* DC3000 *hrcC* as measured by qRT-PCR and normalized against *ACT2*. Samples were collected at different time points after infection. RNA levels are expressed relative to the Col-0 wild-type plants mock treated (B) or infected Col-0 plants (C). Values are from n = 3 biologically independent samples. (D) Quantification of the formation of *Loop^EFR^* and *Loop^Xl-k^*as measured by 3C-qPCR analysis in Col-0 wild-type and *eair-1* plants mock treated or infected for one day with *Pst* DC3000 *hrcC*. Values of n = 4 biologically independent samples are expressed relative to the mock treated samples. In all plots, data are presented as mean values ± s.d. *P* values as calculated with a two-tailed unpaired t-test with Welch’s correction.

Induction of *EFR* was less pronounced in *dcl234* or *eair-1* mutants, indicating that the RdDM pathway and *Ea-IR* are both required for *EFR* upregulation after pathogen infection (Figure 2C). We therefore assessed changes in local chromatin topology using 3C-qPCR assays, which revealed that infection of Col-0 plants with *Pst* DC3000 *hrcC* led to the opening of the *Loop^EFR^* (Figure 2D), in support of a repressive role of the closed loop. In addition, opening of *Loop^EFR^* correlated with closing of *Loop^XI-k^* (Figure 2D), suggesting again that the formation of *Loop^XI-k^* is contingent upon the opening of *Loop^EFR^*. No significant changes in chromatin topology were observed in *eair-1* mutants as could be expected from our result given the lack of an intact *EaIR* (Figure 2D).

### Absence of *Ea-IR* in natural accessions eliminates chromatin loop formation and enhances pathogen resistance

That *Ea-IR* corresponds to a transposon insertion raised the possibility that it might be polymorphic in *A. thaliana* populations and influence pathogen responses in natural accessions. We first consulted published transposon insertion polymorphism data based on short sequencing reads ^48^, which revealed that *Ea-IR* was partially or completely absent in nearly three-quarters of 216 accessions (Figure 3A). Because there are tradeoffs in sensitivity and precision when calling transposon insertion polymorphisms from short reads, we compared the *EFR/Xl-k* region in assembled long-read genomes of 149 accessions ^49,50^. Similar to the short-read data, we found that ∼70% of these accessions lacked *Ea-IR* (Figure 3B). The absence of *Ea-IR* typically coincides with the absence of *LIMPET1*, about 6 kb away, but the sequences between *Ea-IR* and *LIMPET* are present in all accessions (Figure 3B). Recombinants in which only *Ea-IR* or *LIMPET1* is found are rare.

**Figure 3.**
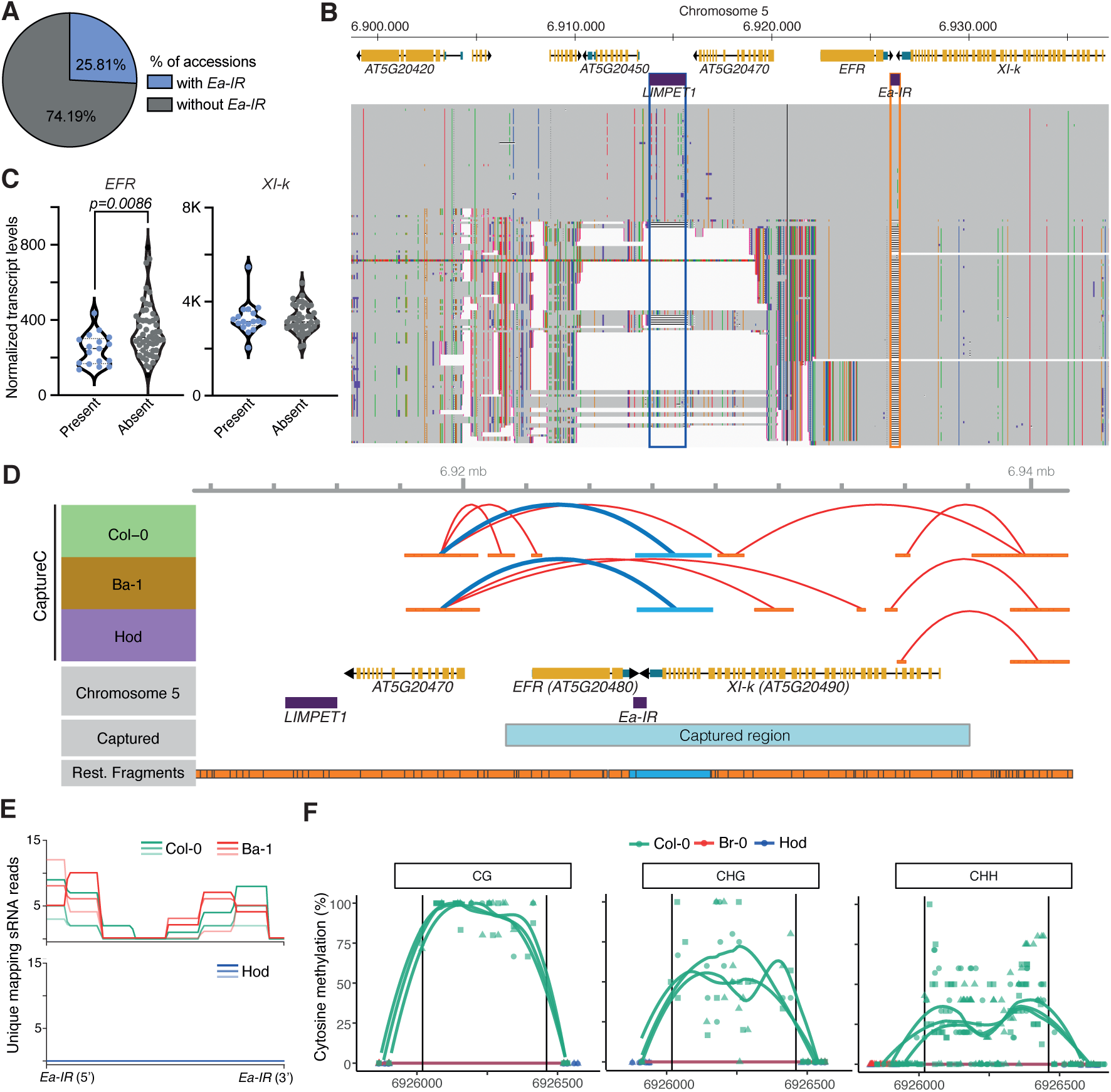
Structural variation at *Ea-IR* in *A. thaliana* natural accessions affects chromatin architecture and *EFR* expression. (A) *Ea-IR* variation, as deduced from a published short-read based variant atlas in 216 natural accessions from the 1001 Genomes Project. (B) Multiple sequence alignment of *EFR/Xl-k* region in genomes assembled from long reads. Each line represents one genome. Single nucleotide polymorphisms are indicated by short vertical-colored lines, relatively small deletions are indicated as horizontal black lines and large deletions are indicated as white space. *LIMPET1* and *Ea-IR* are highlighted with blue and orange lines. The genomes are ordered based on the presence or absence of the *Ea-IR* as mapped to the Col-0 reference genome. (C) Expression of EFR and XI-k in 216 accessions, as measured by RNA-seq from ^88^. Each dot represents the normalized expression value for accessions with or without Ea-IR. *P* value was calculated with a two-tailed unpaired t-test with Welch’s correction. (D) Chromatin contacts from CaptureC in the *EFR/Xl-k* region in Col-0, Ba-1, and Hod accessions. Color code as in Fig. 1D. (E) 24 nt siRNAs mapping uniquely over *Ea-IR* in Col-0, Ba-1, and Hod accessions. Three individual sRNA-seq replicates are plotted separately and shown with different color tones. (F) Cytosine methylation in the CG, CHG, and CG contexts at *Ea-IR* (demarcated by vertical black lines) in Col-0, Br-0, and Hod accessions. Three individual BS-seq replicates are plotted for each genotype.

There was a clear association between the absence of *Ea-IR* and elevated *EFR* expression levels in most accessions, while *XI-k* expression was unaffected (Figure 3C). This result again highlights the dependency of *EFR* expression, as opposed to *XI-k*, on *Ea-IR*. Likewise, when comparing our CaptureC data between two accessions with *Ea-IR* (Col-0 and Ba-1) and one lacking *Ea-IR* (Hod), we observed that *Loop^EFR^* was only formed in Col-0 and Ba-1, but not in Hod (Figure 3D). As anticipated, siRNAs and cytosine methylation associated with *Ea-IR* in Col-0 and Ba-1 were absent in Hod and Br-0, which also lack the IR (Figures 3E and 3F). These data again support the idea that *Ea-IR* plays a critical role in chromatin looping and *EFR* expression.

To further assess the influence of structural variation at *Ea-IR* on *Loop^EFR^* formation and *EFR* expression, we first confirmed that we could detect *Ea-IR* status in the four selected polymorphic accessions using PCR (Figure S1B and S2). We then profiled and quantified the formation of *Loop^EFR^*and *Loop^XI-k^* using 3C-qPCR, finding that the presence of *Ea-IR* is essential for establishing *Loop^EFR^* but dispensable for *Loop^XI-k^*, formed by a region that was present in all four accessions (Figure 4A, mock-treated samples). Nevertheless, as previously observed, the absence of *Loop^EFR^*in Hod and Br-0 -caused by the lack of *Ea-IR* enhanced the formation of *Loop^XI-k^* in these plants (Figure 4A, mock-treated samples). The accessions with *Ea-IR* had markedly lower *EFR* expression levels, in line with the hypothesized repressive role of *Loop^EFR^* (Figure 4B, mock-treated samples). In contrast, the levels of *XI-k*, which we had determined to be independent of *Ea-IR* in Col-0, were similar among accessions (Figure 4B, mock-treated samples).

**Figure 4.**
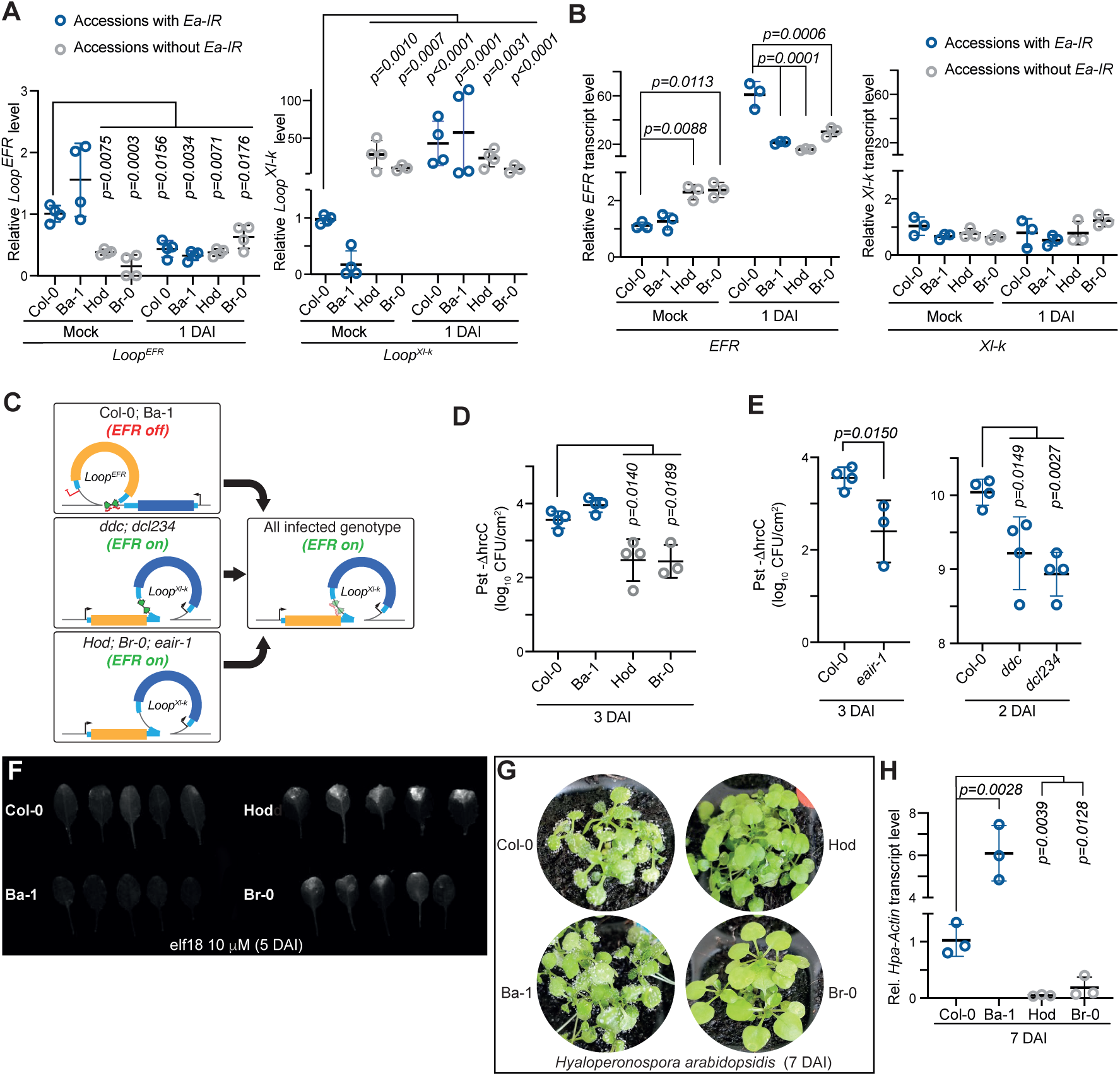
Polymorphic accessions lacking the *Ea-IR* present heightened resistance to pathogens. (A) Quantification of *Loop^EFR^* and *Loop^Xl-k^* as measured by 3C-qPCR in Col-0, Ba-1, Hod, and Br-0 accessions infected with *Pst* DC3000 *hrcC* (1 dai). Values of n = 4 biologically independent samples are expressed relative to the mock-treated samples. Data are presented as mean values ± s.d. *P* values were calculated with a two-tailed unpaired t-test with Welch’s correction. (B) *EFR* (left) and *Xl-k* (right) expression in the same samples as in (A). RNA levels are expressed relative to the mock-treated Col-0 wild-type plants. Values of n = 3 biologically independent samples. Data are presented as mean values ± s.d. *P* values were calculated with a two-tailed unpaired t-test with Welch’s correction. (C) Diagram of *Ea-IR* controlled chromatin dynamics in different backgrounds. *EFR* gene model is yellow, *Xl-k* blue, and *Ea-IR* is indicated by green arrows. Red pins represent cytosine methylation. Expression status of *EFR* (“on”/“off”) is indicated in green. Left, before infection. Right, after infection, where all backgrounds converge on a similar chromatin organization. (D and E) Growth of *Pst* DC3000 *hrcC* in accessions (D) or mutants (E), expressed as the log_10_ of the colony forming units per cm^2^ of infected tissue. Values of n = 4 biologically independent samples. Data are presented as mean values ± s.d. *P* values were calculated with a two-tailed unpaired t-test with Welch’s correction. (F) Increased autofluorescence of *A. thaliana* leaves, indicative of cell death, after treatment with elf18 peptide in accessions with *Ea-IR* (Col-0 and Ba-1). (G and H) Increased susceptibility to *Hyaloperonospora arabidopsidis* (*Hpa*) in accessions with *Ea-IR* (Col-0 and Ba-1), as observed by sporulation 7 dai (G) or as quantified by RT-qPCR of *Hpa* actin gene expression, normalized to host *ACT2*. Values of n = 3 biologically independent samples. Data are presented as mean values ± s.d. *P* values were calculated with a two-tailed unpaired t-test with Welch’s correction.

Our next inquiry was whether structural variation at *Ea-IR* alters the effects of *Pst* DC3000 *hrcC* infection on chromatin organization and *EFR* expression. Like Col-0, infection led to the opening of *Loop^EFR^* and the concurrent emergence of *Loop^XI-k^* in Ba-1 (Figure 4A, 1 DAI). Unlike the situation in Col-0 and Ba-1, infection affected neither *Loop^EFR^*nor *Loop^XI-k^* in the accessions lacking *Ea-IR* (Figure 4A, 1 DAI). Concordantly, Hod and Br-0 had substantially higher levels of *EFR* expression even before infection. Equivalent levels were reached in the accessions with *Ea-IR* only late during infection (Figure 4B), when both chromatin organization and *EFR* expression levels had become very similar in all four tested accessions (Figure 4A and B). The role of *Ea-IR* can thus be interpreted as reducing *EFR* expression in unchallenged plants. That *Ea-IR* is relatively widespread in natural *A. thaliana* populations suggests a considerable selective advantage of reduced *EFR* expression in the absence of infection, at least in some environments (summarized in Figure 4C).

To test whether the differences in chromatin and *EFR* expression dynamics translate into varying pathogen responses, we challenged the four accessions along with *ddc*, *dcl234*, and *eair-1* mutants with *Pst* DC3000 *hrcC* and monitored bacterial growth. In line with the higher basal levels of the *EFR* immune receptor, the three mutants and the two accessions without *Ea-IR*, Hod, and Br-0, supported much less pathogen growth than accessions with *Ea-IR* (Figure 4D and 4E). To directly test the role of *EFR*, we treated the four accessions with 10 μM of elf18, an *EFR* ligand, and measured autofluorescence as an indicator of cell death after five days. Signs of cell death were much more prominent in the two accessions without *Ea-IR*, indicating the increased basal levels of *EFR* expression are protective and confirming that the higher ERF expression levels translate into more active receptors (Figure 4F). Lack of *Ea-IR* also increased resistance to the oomycete *Hyaloperonospora arabidopsidis* (*Hpa*) (Figure 4G and 4H). Since *Hpa* does not produce EFR ligands, we conclude that increased resistance is due to the higher background levels of *EFR* in the accessions without *Ea-IR*, likely causing mild constitutive immunity.

### Opening of *Loop^EFR^* leads to Pol II readthrough and generation of an *XI-k* transcript containing *Ea-IR* sequences

Pol II commonly transcribes regulatory inverted repeats (IRs) within 500 bp of genes, generating 24 nucleotide siRNAs via a non-canonical DCL3-dependent pathway ^34,38^. There are at least eight other copies of *Ea-IR* like sequences across the Col-0 genome (Figure S4A), all near protein-coding genes, and most being found in other accessions (Figure S4B and S4C). However, there are uniquely mapping siRNA that come from the *Ea-IR* locus between *EFR* and *Xl-k,* indicating that this element is indeed transcribed (Figure S4D).

To understand how *Ea-IR* is transcribed, we analyzed Pol IV and Pol V chromatin immunoprecipitation followed by sequencing (ChIP-seq) data alongside Pol II native elongating transcript sequencing (NET-seq) data. As with other transposons, the ChIP-seq data indicated that Pol IV and Pol V likely contribute to *Ea-IR* transcription (Figure 5A), but the NET-seq data revealed that Pol II can also transcribe this region. The specific distribution of NET-seq reads suggested that Pol II-dependent *Ea-IR*-transcripts contain also *XI-k* mRNA sequences (Figure 5A). In animals, specific stressful conditions can trigger Pol II to read through termination sites, producing transcripts that include sequences from nearby transposons ^51-53^. Similarly, we observed Pol II readthrough at the termination sites of *XI-k*, leading to the incorporation of *Ea-IR* sequences into *Xl-k* transcripts (Figure 5A). Specific primers allowed us to identify and quantify the extended *XI-k* transcript that includes *Ea-IR* sequences (‘*XI-k* long transcript’, Figure S5). Long *XI-k* transcript levels are very low in Col-0 plants, but increase substantially upon pathogen infection (Figure 5B and 5C). This observation potentially suggests that *Loop^EFR^*, which opens after infection, impedes Pol II readthrough. To test this hypothesis, we measured the levels of the long *XI-k* transcript in *ddc* and *dcl234* mutants with a constitutively open *Loop^EFR^*. Long *XI-k* transcript levels were significantly increased in both mutants (Figure 5D).

**Figure 5.**
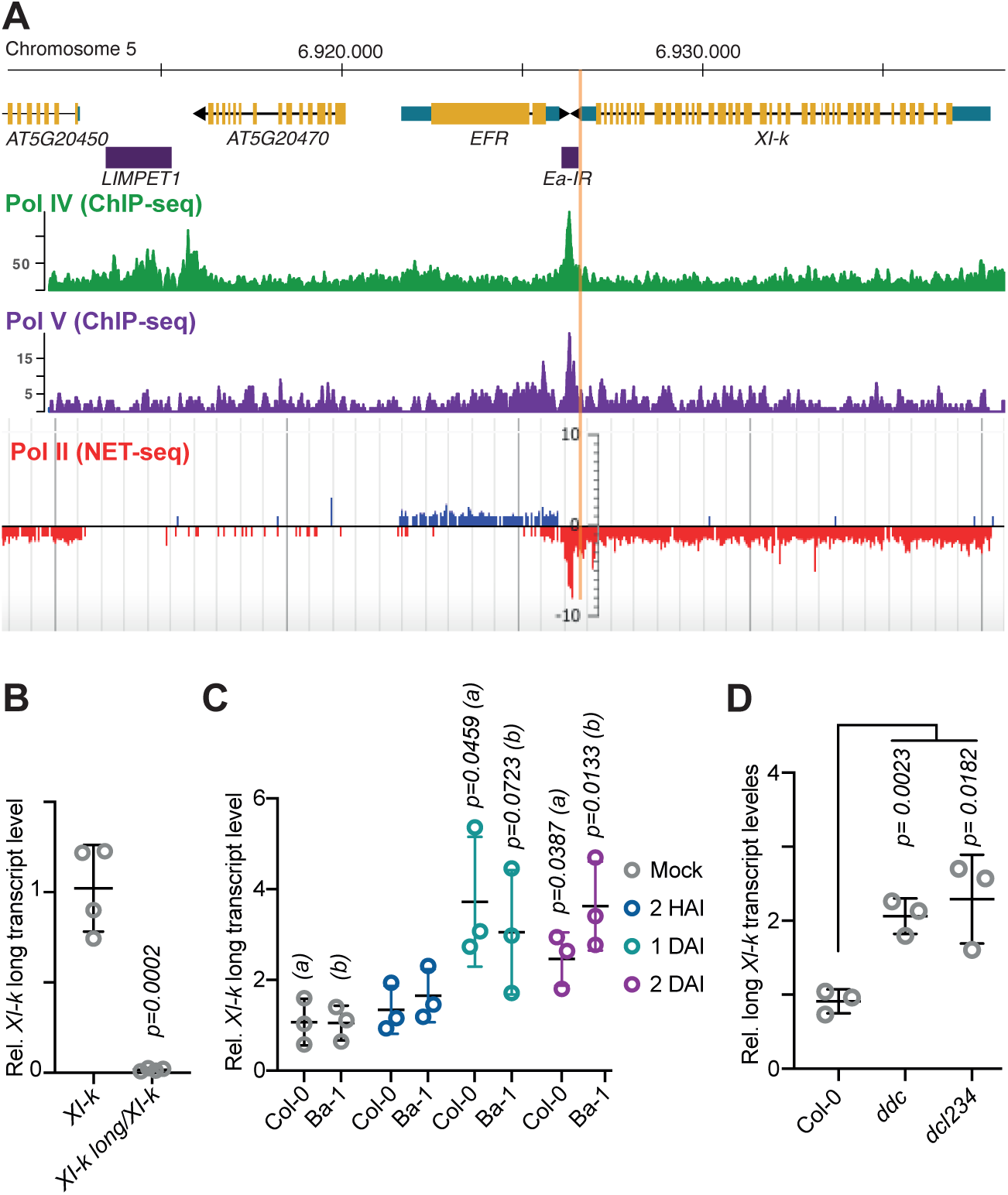
Pathogen-induced opening of *Loop^EFR^* leads to Pol II readthrough at *Xl-k* into *Ea-IR*. (A) Integrated Genomics Viewer (IGV) genome browser views of Pol IV and Pol V ChIP-seq ^89^ and Pol II nascent elongating transcripts ^90^ in untreated Col-0 wild-type plants. Scale is normalized to total mapped reads. Note that Pol II signal extends beyond the *Xl-k* termination site, indicated by a vertical orange line. (B, C and D) Expression of the canonical and the long *Xl-k* transcript isoforms, as measured by qRT-PCR in untreated Col-0 wild-type plants (B); in accessions with *Ea-IR* (Col-0 and Ba-1) after different time points of infection with *Pst* DC3000 *hrcC* (C); and in Col-0, *ddc,* and*dcl234* mutants (D). Values of n = 4 (B) and 3 (C and D) biologically independent samples are expressed relative to the Col-0 plants. Data are presented as mean values ± s.d. *P* values were calculated with a two-tailed unpaired t-test with Welch’s correction.

### Long *Xl-k* transcripts with *Ea-IR* sequences are processed by DCL1 and DCL3 to produce siRNAs

RNA hairpin structures, as in the long *Xl-k* transcript, can not only affect mRNA transcription and translation but also serve as substrates for DICER/DROSHA-mediated processing into small RNAs ^54-58^. Consistent with this last notion, we found that the abundance of the long *XI-k* transcript is much higher in *dcl1-100* mutants (Figure 6A), indicating that this transcript with *Ea-IR* sequences is subject to DCL1-dependent processing and degradation. To test this directly, we transiently expressed in *Nicotiana benthamiana* chimeric transcripts that included GFP coding sequences and *Ea-IR* sequences. Inserting *Ea-IR* downstream of GFP, but not upstream of GFP, reduced GFP fluorescence, suggesting either transcript degradation or lack of translation (Figure 6B). The simultaneous introduction of an artificial miRNA silencing DCL1 or its HYL1 cofactor restored GFP fluorescence levels to those of controls, confirming that the reduction in GFP signal was indeed DCL1-dependent (Figure 6B). The 21 nt siRNAs derived from *Ea-IR* were also produced in *rdr6* mutants and in *pol-IV* mutants (Figure 6C). Their independence of the RDR6/DCL4 pathway differentiates them from the 21 nt siRNAs originating from the nearby *LIMPET1* transposons, which behaves in this respect as other canonical transposons, where the dsRNA substrate for DCL processing into 21 nt sRNA does not come from a self-complementary hairpin structure but requires the copying of the primary transcript by RDR6 (Figure 6C, ^59^. Levels of 21 nt *Ea-IR*-derived siRNAs are particularly high in *dcl3* mutants, with a concurrent decrease in 24 nt siRNAs (Figure 6C), suggesting that processing of *Ea-IR* transcripts involves competition between DCL1 and DCL3. We introduced the *GFP-Ea-IR* and *Ea-IR-GFP* constructs into *dcl234* mutants, and found that GFP fluorescence of both was much higher in the mutant plants, indicating that DCLs other than DCL1, most likely DCL3 as suggested by the drop in 24 nt sRNAs, contribute to processing of the long *XI-k* transcript containing *Ea-IR* sequences (Figure 6D).

**Figure 6.**
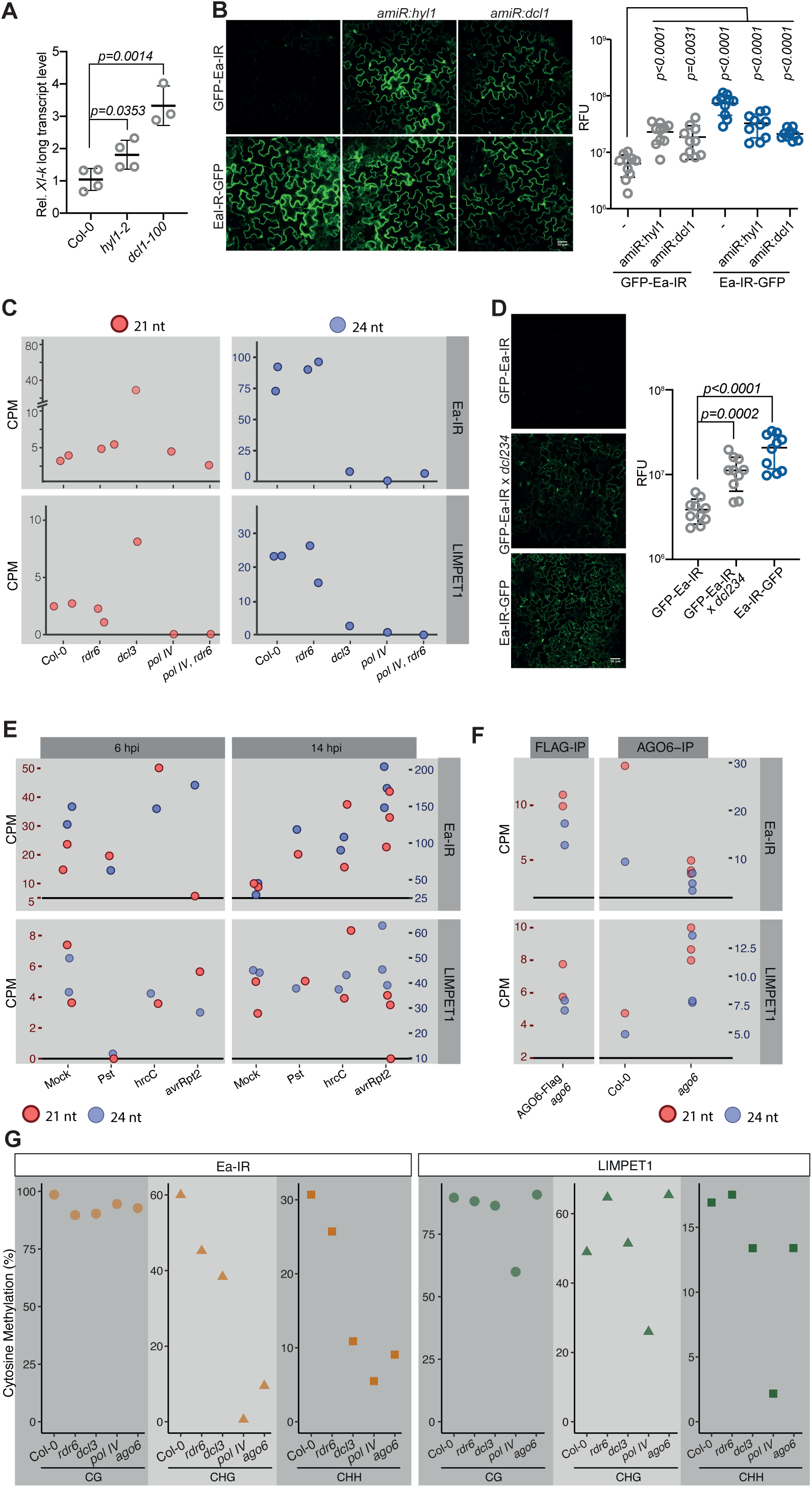
DCL1 and DCL3 process the *Ea-IR*-containing long *XI-k* transcript isoform to produce epigenetically active sRNAs. (A) Expression of the canonical and the long *Xl-k* transcript isoform in untreated Col-0 wild-type, *hyl1-2* and *dcl1-100* plants. Values of n = 4 biologically independent samples are expressed relative to the Col-0 plants. Data are presented as mean values ± s.d. *P* values were calculated with a two-tailed unpaired t-test with Welch’s correction. (B) Left, Confocal microscopy of *Nicotiana benthamiana* leaves expressing constructs with the *Ea-IR* sequence inserted upstream or downstream of the GFP coding sequence (Ea-IR-GFP and GFP-Ea-IR). Leaves were co-infiltrated with an empty vector or artificial miRNAs targeting *HYL1* and *DCL1*. Scale bars 50 μm. Right, quantification of fluorescence intensity of leaves expressing from the left panel. Data are presented as mean values ± s.d. of 10 independent leaves. *P* values were calculated with a two-tailed unpaired t-test with Welch’s correction. (C) 21 nt and 24 nt sRNAs (cpm, counts per million) ^86^ mapping to the *Ea-IR* and *LIMPET1* transposons in Col-0 wild-type, and *rdr6, dcl3, pol IV*, and *pol IV rdr6* plants. (D) Confocal microscopy images and quantification of GFP-Ea-IR fluorescence in transgenic *A. thaliana* wild-type and *dcl234* plants. Transgenic plants expressing Ea-IR-GFP were included as a positive control. Scale bars 50 μm. Data are presented as mean values ± s.d. of 10 independent leaves. P values were calculated with a two-tailed unpaired t-test with Welch’s correction. (E) 21 nt and 24 nt sRNAs (cpm, counts per million) ^87^ mapping to the *Ea-IR* and *LIMPET1* transposons in Col-0 wild-type samples 6 or 14 hours after infection with a virulent (*Pst* DC3000), a type III secretion system-deficient (*Pst* DC3000 *hrcC*), or an avirulent (*Pst* DC3000 avrRpt2) *P. syringae* strain. (F) 21 nt (red dots) and 24 nt (blue dots) sRNAs associated with the endogenous or a FLAG-tagged version of AGO6 ^64^ mapping to the *Ea-IR* and *LIMPET1* transposons in Col-0 wild-type plants, *ago6* mutants, and *AGO6-FLAG ago6* complemented transgenic plants. (G) Percentage of cytosine methylation in the CG, CHG, and CHH context mapping the *Ea-IR* and *LIMPET1* transposons in Col-0 wild type plants and *rdr6*, *dcl3*, *pol IV*, and *ago6* mutant plants. Each dot shows the average cytosine methylation over the selected region as detected by BS-sequencing experiments^86^_.._

So far, our findings showed that pathogen infection causes Pol II readthrough at *XI-k* and generation of transcripts that contain *Ea-IR* sequences, which can undergo DCL1-mediated processing into siRNAs. We confirmed that pathogen challenge causes an increase in 21 nt *Ea-IR*-derived siRNAs as early as 14 hours after infection (Figure 6E). This induction is specific and not seen at the nearby *LIMPET1* transposon. This observation is in agreement with the opening of *Loop^EFR^* during infection, the increase in the long *XI-k* transcript and DCL1-dependent processing of this transcript isoform into siRNAs. All tested *P. syringae* strains (*Pst* DC3000, *Pst* DC3000 *hrcC*, and *Pst* DC3000 *avrRpt2*) induce the production of *Ea-IR*-derived 21 nt siRNAs (Figure 6E), suggesting that the signal leading to the production of the long *Xl-k* transcript isoform is likely a PAMP signal.

The action of the RdDM pathway can be initiated by 21 nt siRNAs by facilitating the recruitment of Pol IV and Pol V ^59-63^. Such siRNAs are predominantly loaded into AGO6 ^64^, and this is also the case for *Ea-IR*-derived 21 nt siRNAs (Figure 6F, ^64^). Supporting the scenario of DCL1-dependent RdDM initiation, we observed CHH and CHG methylation over *Ea-IR* in *dcl3*, *rdr6*, and *pol IV* mutants (Figure 6F). Once initiated, methylation maintenance likely relies on the canonical DNA methylation pathway, as indicated by the presence of Pol IV and Pol V associated with *Ea-IR* (Figure 5A).

## Discussion

Plants are constantly exposed to a diverse range of pathogens in their environment. Swift activation of defense responses is therefore vital for plant survival but it must be balanced with the risk of an overactive immune system, which can negatively affect growth and yield ^65,66^. Conversely, an inadequate immune response leaves plants vulnerable to pathogens ^24,25,65,67^. Our study discloses a new regulatory mechanism for the activation and deactivation of immune responses, with epigenetic control of the expression of the *EFR* immune receptor gene. We discovered that a transposon-derived IR, *Ea-IR*, located downstream of *EFR*, orchestrates the formation of a repressive chromatin loop that suppresses *EFR* transcription in plants not yet exposed to pathogens. That such a dampened immune state is often not advantageous can be inferred from the finding that *Ea-IR* is found only in a subset of *A. thaliana* natural accessions. Plants without *Ea-IR* had higher basal activity of the immune system, which is likely beneficial in a pathogen-rich environment.

Activation of EFR by fragments of bacterial elongation factor Tu (EF-Tu) requires the co-receptor BRI1-ASSOCIATED RECEPTOR KINASE 1 (BAK1). Because BAK1 is also involved in brassinosteroid signaling ^2,13^, competition for this rate-limiting co-receptor contributes to defense-growth trade-offs ^25,68^. Importantly, *Ea-IR* not only limits *EFR* expression before acute exposure to pathogens but also helps to dampen *EFR* expression after its induction. During infection, the open *Loop^EFR^* state not only leads to increased *EFR* expression, but also allows Pol II readthrough at the neighboring *Xl-k* gene. The long *Xl-k* transcript isoform generated by readthrough includes *Ea-IR* sequences, which can be processed by DCL1 or DCL3 into epigenetically active siRNAs. These siRNAs can then trigger the methylation of *Ea-IR*, restoring *Loop^EFR^* and *EFR* repression (Figure 7).

**Figure 7.**
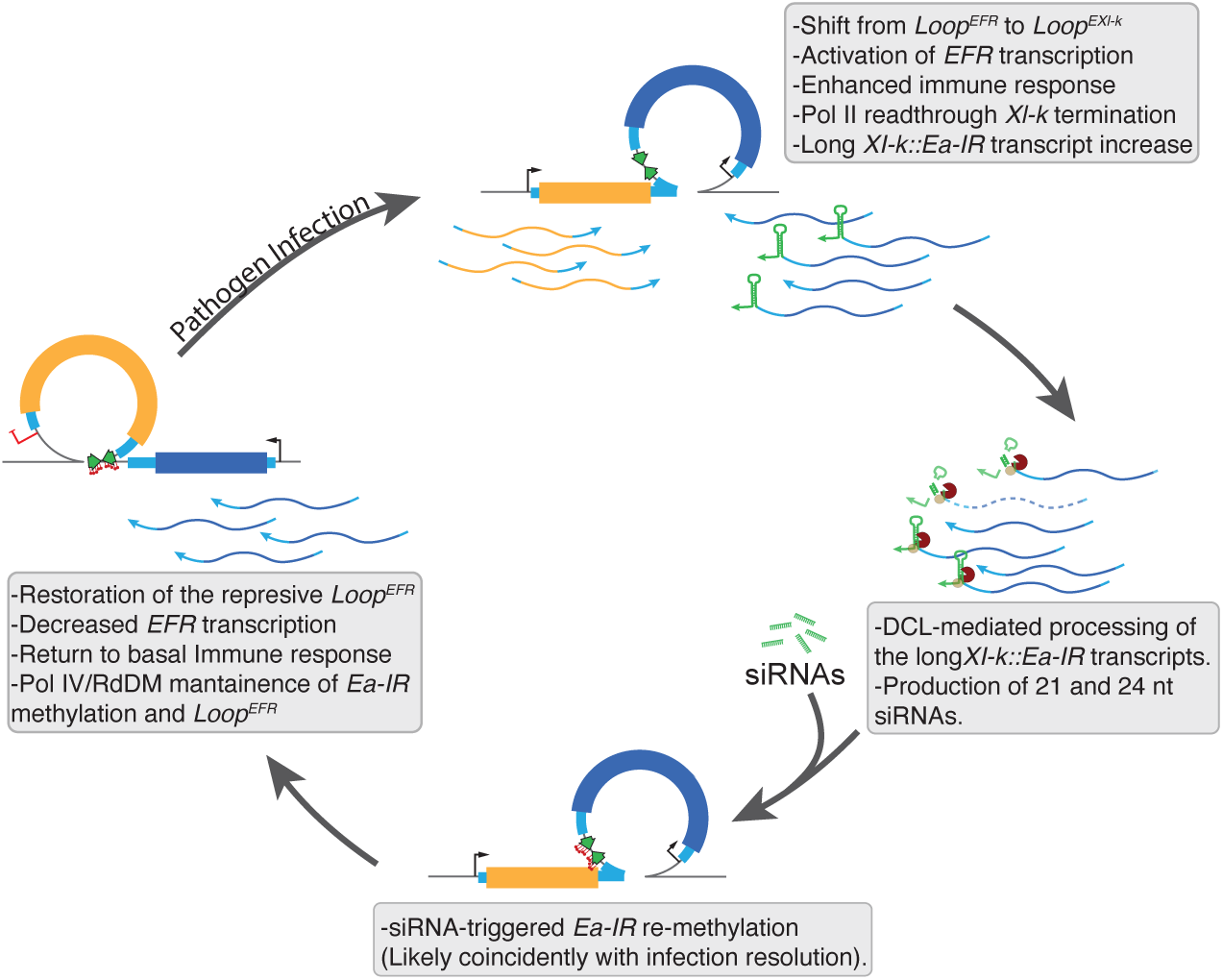
A proposed model for a self-buffered immune response mechanism controlled by the *Ea-IR*-mediated regulation of chromatin organization and transcription at EFR locus. In a resting state, *A. thaliana* plants with the *Ea-IR* insertion between *EFR* and *Xl-k* genes from the repressive *Loop^EFR^* chromatin loop, which encompasses *EFR* coding sequences. This loop dampens the background activity of the immune system and also promotes the termination of *Xl-k* transcription. Upon pathogen infection, the repressive *Loop^EFR^* opens, allowing for enhanced *EFR* transcription, which increases immunity and allowing Pol II readthrough at the *Xl-k* termination site. The hairpin at the 3’-end of the long *Xl-k* transcript isoform is processed by DCL1 and DCL3 into epigenetically active siRNAs. These siRNAs can subsequently initiate the re-methylation of *Ea-IR* sequences, promoting the reformation of *Loop^EFR^* and thereby resetting chromatin topology to a homeostatic state with a steady but reduced immune response. And open *Loop^EFR^*or the absence of *Ea-IR* allows for the formation of a second, apparently innocuous *Loop^Xl-k^*.

Receptor attenuation is a well-known mechanism in biology that involves the downregulation or reduction in the activity of cell surface receptors in response to various stimuli. Typically, this process occurs at the post-translational level, where receptors are either internalized, degraded, or otherwise modified to reduce their signaling capacity. Interestingly, our discoveries share some similarities with receptor attenuation, but at a transcriptional level. In our case, the increment of EFR during pathogen infection is quickly attenuated once the infection is resolved by re-establishing the repressive chromatin loop and reducing EFR receptor expression. This mechanism resembles receptor attenuation in that it modulates receptor levels but does so by controlling transcription rather than post-translational modifications.

As a consequence of the high level of conservation of EF-Tu protein sequences across bacteria, increased levels of EFR confer resistance against a wide range of bacterial pathogens ^69^. Conversely, transgenic plants expressing EFR do not show enhanced resistance to the fungal pathogen *Verticillium dahliae* ^69^. Our data showed that the lack of *Ea-IR* in natural accessions increased resistance to the oomycete *Hpa* (Figure 4G and 4H). Such difference could be the consequence of the necrotrophic vs biotrophic pathogens nature of both pathogens. this goes in line with the report that BAK1, an EFR/FLS2 shared co-receptor, contribute to disease resistance against the hemibiotrophic bacterium *Pst* and the obligate biotrophic oomycete *Hpa* ^70^. Possibly, such resistance is achieved by the constitutively higher levels of EFR in *Ea-IR*-missing accessions that could produce a primed state of the immune system through the constant interaction of plants to non-pathogenic bacteria still recognized by EFR. While EFR-induced defense may be constitutively higher in plants missing *Ea-IR*, a TE-induced priming of the defense through de-repression of the EFR locus is possible in those accessions with the IR in place. TE-induced priming of immune response has been previously reported ^71-74^. Whether epigenetic changes in chromatin organization at *EFR* locus could lead to a primmed state remains to be explored.

Since *Hpa* does not produce EFR ligands, we conclude that increased resistance is due to the higher background levels of *EFR* in the accessions without *Ea-IR*, likely causing mild constitutive immunity.

Epigenetically active siRNAs of 21 nt, especially when loaded in AGO6, can initiate RdDM ^63,64^. While canonical RdDM pathways, led by DCL3-dependent 24 nt siRNAs, may initiate *Ea-IR* DNA methylation after infection, DCL1-dependent 21 nt sRNAs, produced from the long *Xl-k* transcript, might also participate in this process. The association of Pol IV and Pol V with the *Ea-IR* encoding region suggests that the long-term maintenance of DNA methylation, and thus the *Loop^EFR^*, follows a more canonical RdDM pathway. Nevertheless, the presence of additional copies of the *Ea-IR* in the *A. thaliana* genomes opens the possibility of regulatory crosstalk between them to initiate or maintain DNA methylation. Whether the structural nature of the *Ea-IR* transcript is sufficient to trigger the reported regulatory mechanism independently of the specific *Ea-IR* sequence or whether this IR has special properties in terms of recruiting specific elements of the DNA methylation machinery remains to be explored. However, the first scenario appears more likely, as we found that many IRs, regardless of their sequence, promote chromatin reorganization when expressed ^34^. Still, it is possible that some sequence-specific DNA methyl readers can recognize methylated *Ea-IR* and fine-tune its regulatory function. The fact that only the expression of *EFR*, but not *Xl-k*, is regulated by *Ea-IR* argues against a repressor binding the IR to directly regulate the locus.

Transcriptional readthrough, observed in various organisms, might play an important role in small genomes like that of *A. thaliana* where genes are closely spaced. Our data suggest that *Loop^EFR^* formation inhibits readthrough, probably serving as a barrier for Pol II elongation and promoting transcriptional termination. This feature of *Loop^EFR^* unveils a mechanism of action for short-range chromatin loops that has not been described before.

Our study uncovers a complex epigenetic regulatory mechanism where IR-derived siRNAs modulate chromatin topology and *EFR* expression to shape the dynamic of plant immune responses. *Ea-IR* is a fascinating example of a genetic allele that acts through epigenetic changes ^75^, with the modulation of loop formation providing an elegant mechanism for rapid transcriptional responses to pathogens. It also shows that cis-regulatory variation is not restricted to differences in the recruitment of DNA-binding transcription factors. This work shows how the evolutionary loss of specific IRs can determine a plant’s ability to mount an effective immune response, highlighting the importance of chromatin organization in shaping adaptive traits. Understanding these processes also opens avenues for improving crops’ stress resistance and pathogen resilience through manipulating IR elements without altering coding sequences.

## Materials and Methods

### Plant Material and Growth Conditions

*Arabidopsis thaliana* and *Nicotiana benthamiana* plants were cultivated in soil for 3 to 4 weeks under long-day conditions (16 hours of light and 8 hours of darkness). *Arabidopsis thaliana* accessions Col-0, Hod, Ba-1, and Br-0 were used. T-DNA insertion mutants were *dcl2 dcl3 dcl4* (*dcl234*) ^76^, *drm1 drm2 cmt3* (*ddc*) ^77^, *eair-1* (WiscDsLox440E09), *hyl1-2* (SALK_064863), and *dcl1-100* (GABI_098F10). Transgenic plants contained the constructs *35S::GFP-Ea-IR* and *35S::Ea-IR-GFP*, for which and *Ea-IR* fragment was amplified using Phusion polymerase (Thermo Fisher), then cloned into pEntr/D-TOPO (Thermo Fisher), and subsequently recombined using LR Clonase (Thermo Fisher) into Gateway-compatible pGreen destination vectors (pCB012 and pCB011, respectively). *Arabidopsis thaliana* plants were transformed using the floral dip method.

### Transient Expression in *Nicotiana benthamiana*

For transient expression in *Nicotiana benthamiana*, *Agrobacterium tumefaciens* GV3101 carrying GFP-Ea-IR, Ea-IR-GFP, *amiR:HYL1*, or *amiR:DCL1* constructs were cultured for 20 hours from frozen stocks on LB medium supplemented with antibiotics. The bacterial cultures were then resuspended in infiltration medium (containing 10 mM MgCl2, 10 mM MES [pH 5.7], and 150 μM acetosyringone) at an OD_600_ of 0.5 and incubated for 3 hours at room temperature. Subsequently, plants were infiltrated using equal parts of these bacterial suspensions. Samples were analyzed after 2 days. Artificial miRNA constructs were designed following the methodology outlined by ^78^.

### Comparison of *EFR/Xl-k* region in *A. thaliana* genomes

A set of 149 *Arabidopsis thaliana* accessions genomes sequenced with the PacBio technology ^49,50^ were analyzed. The obtained long reads were mapped to the TAIR10 genome sequence with minimap2, version 2.24 ^79^; using the -ax asm5 parameter. The alignments on the EFR/XI-k region were then inspected with the Integrative Genomics Viewer ^80^. Expression of *EFR* and *XI-k* in 216 accessions is expressed as normalized gene expression levels from datasets retrieved from NCBI’s Gene Expression Omnibus, with the GSE80744 series identifier (file GSE80744_ath1001_tx_norm_2016-04-21-UQ_gNorm_normCounts_k4.tsv). Accessions included correspond to those analyzed by ^48^.

### Detection of *Ea-IR* Presence and T-DNA Insertions

IR presence and T-DNA insertions were assessed through PCR using DreamTaq polymerase (ThermoFisher) with genomic DNA. For genomic DNA extraction, 100 mg of fresh plant material was ground in 700 μl of extraction buffer (comprising 200 mM Tris-HCl pH 8, 250 mM EDTA, 0.5% SDS), followed by precipitation with isopropanol. PCR was conducted using primers as detailed in Table S1. Molecular weight marker GeneRuler 1 kb plus (ThermoFisher) was used to infer band size.

### 3C Assay and RT-PCR

The 3C assay was conducted following a published procedure ^33^. Approximately 2 g of tissue was collected per sample and crosslinked with 1% formaldehyde. Crosslinking was stopped by adding glycine to a final concentration of 0.125 M, after which nuclei were isolated.

To detect loops 1 to 8, including *Loop^EFR^* and *Loop^XI-k^*, DNA was digested with XbaI (ThermoFisher). For loops 9 and 10, Bsp1407I (ThermoFisher) was used, and for loop 11, EcoRI (ThermoFisher). DNA fragments were ligated with 100 U of highly concentrated T4 DNA ligase (Thermo) at 22°C for 5 hours in a 2 ml volume. Subsequent steps included reverse crosslinking, proteinase K treatment (QIAGEN), and DNA purification with the phenol/chloroform method. To measure interaction frequency, qPCR was performed, and results were analyzed with the 2^−ΔΔCt^ formula, using *ACTIN2* as a control. Oligonucleotide primers are given in Table S1. The final qPCR products were subjected to agarose gel electrophoresis (2%), and bands corresponding to the expected size were purified and sequenced using the Sanger method to confirm ligation products.

For quantitative RT-PCR, 500 ng of total RNA underwent reverse transcription using the RevertAid RT Reverse Transcription Kit (ThermoFisher) after treatment with DNase (ThermoFisher). Subsequently, qPCR was conducted using SYBR green (ThermoFisher Maxima SYBR Green qPCR Master Mix (2x)). Relative expression levels were determined using the 2^−ΔΔCt^ method, with *ACTIN2* as the housekeeping gene control. Oligonucleotide primers are given in Table S1.

### Plant Inoculation and Pathogen Growth Assays

To assess disease phenotypes, gene expression, and chromatin conformation, plants were subjected to syringe infiltration with the pathogen *Pseudomonas syringae* pv. *tomato* strain DC300 *hrcC* (*Pst* DC3000 *hrcC*), following the method described in ^81^. For this purpose, bacteria were streaked from frozen stocks onto King’s B medium supplemented with Rifampicin. After transferring the bacteria into liquid King’s B medium, they were grown at 28°C for 4 hours and then resuspended at an OD_600_ of 0.002 in sterile water. Fully developed leaves were infiltrated on the abaxial side using a needleless syringe. In cases where assays involved *drm1 drm2 cmt3* (*ddc*) plants, inoculation was carried out via spray using a bacterial suspension with an OD_600_ of 0.2 due to leaf size constraints. To determine bacterial growth, samples were collected at various time points. Leaf discs were extracted from 8 distinct leaves on separate plants, divided into four replicates, ground, resuspended in 1 ml sterile water, diluted, and plated on agar-King’s B plates supplemented with Rifampicin for colony-forming unit (CFU) counting. Alternatively, in assays involving *ddc* plants, samples were taken by weight.

For elf18 (Proteogenix) treatment, plant leaves were infiltrated on the abaxial side using a needleless syringe with a 10 μM peptide solution in sterile water. After 5 days, cell death around the infiltration area was observed using a UV transilluminator.

In infection assays using *Hyaloperonospora arabidopsidis*, 10-day-old Arabidopsis plants grown in soil were subjected to spray inoculation with a conidiospore density of 105 spores/ml. Post-inoculation, the plants were cultivated under short-day conditions (8 hours of light and 16 hours of darkness) with elevated humidity.

### Confocal Microscopy

Samples were excited at 480 nm to detect GFP, and emission was collected within 503 to 531 nm. All assays were conducted using a TCS SP8 confocal microscope (Leica, Solms, Germany). RFU were calculated using Fiji Software.

### Bisulfite Sequencing

DNA was extracted with the DNeasy Plant Mini Kit (QIAGEN). Subsequently, DNA was fragmented to a size of 350 bp through Covaris ultrasonication. Libraries were constructed with the Illumina TruSeq DNA Nano Kit. After the incorporation of adaptors, libraries underwent bisulfite conversion using the EpiTect Plus DNA Bisulfite Conversion Kit (QIAGEN), followed by treatment with Kapa Hifi Uracil+ DNA polymerase (Kapa Biosystems, USA). Paired-end reads (2 x 150 bp) were on a HiSeq 3000 instrument (Illumina). Sequencing reads were first quality trimmed and filtered with Trimmomatic (version 0.36, ^82^), and Bismark ^83^ with the Bowtie2 mapper was used for quantifying cytosine methylation. The results were analyzed in R (R core team, 2022) with the methylKit package. Only cytosines with a coverage of 4 or more were considered. Chop-qPCR assays were performed using 200 ng of restriction enzyme-digested ("chopped") genomic DNA with 10U of MspI (Methylation-sensitive), as described in ^33^. Uncut genomic DNA was used as a control.

### Small RNA (sRNA) Sequencing and ChIP-Seq analysis

The sRNA-seq data for Col-0, *dcl234,* and *ddc* genotypes were generated and analyzed as described ^34,84^. The sRNA coverage over the *Ea-IR* element was generated with deeptools ^85^, using the computeMatrix and plotProfile commands. Publicly available sRNA-Seq reads were retrieved and analyzed. The sequencing data used were from ^86^ (GSE79780), ^87^ (GSE19694), and ^64^ (GSE57191). ChIP-Seq data for Pol IV (PRJNA193689) and Pol V (PRJNA169238) were used in this study. Reads were quality-trimmed with Trimmomatic and mapped to the genome with STAR. The coverage over the *Ea-IR* locus was visualized with Gviz.

## Acknowledgments

We thank Nicolás Cecchini for sharing the *Pst* DC3000 *hrcC* strain and for fruitful discussions and Matias Capela for critical reading of the manuscript. This work was supported by grants from Agencia Nacional de Promoción Científica y Tecnológica (PICT-2021-I-A-00452), Universidad Nacional del Litoral (CAI+D 2020), and Spanish Ministry of Science and Innovation (PID2022-137037NB-I00) to P.A.M., and by DFG-TRR356 and the Max Planck Society to D.W. P.A.M., A.L.A., and I.T. are members of CONICET; R.M. is a fellow of CONICET. P.A.M is a member of CSIC. We thank the Secretaria de Ciencia de la Provincia de Santa Fe and Company of Biologists for a short-term fellowship to R.M.

## Author Contributions

R.M. performed all wet-biology experiments with the help of A.C., G.S. and C.H. A.L.A. performed the informatic analysis with the help of W.X. R.M., A.L.A., P.A.M. and D.W. conceived the study; P.A.M. and D.W. supervised the work and secured project funding; R.M., D.W. and P.A.M. wrote the manuscript with help from all authors.

