## Supplemental information for "Transposon-triggered epigenetic chromatin dynamics modulate EFR-related pathogen response"

### Supplemental Figures

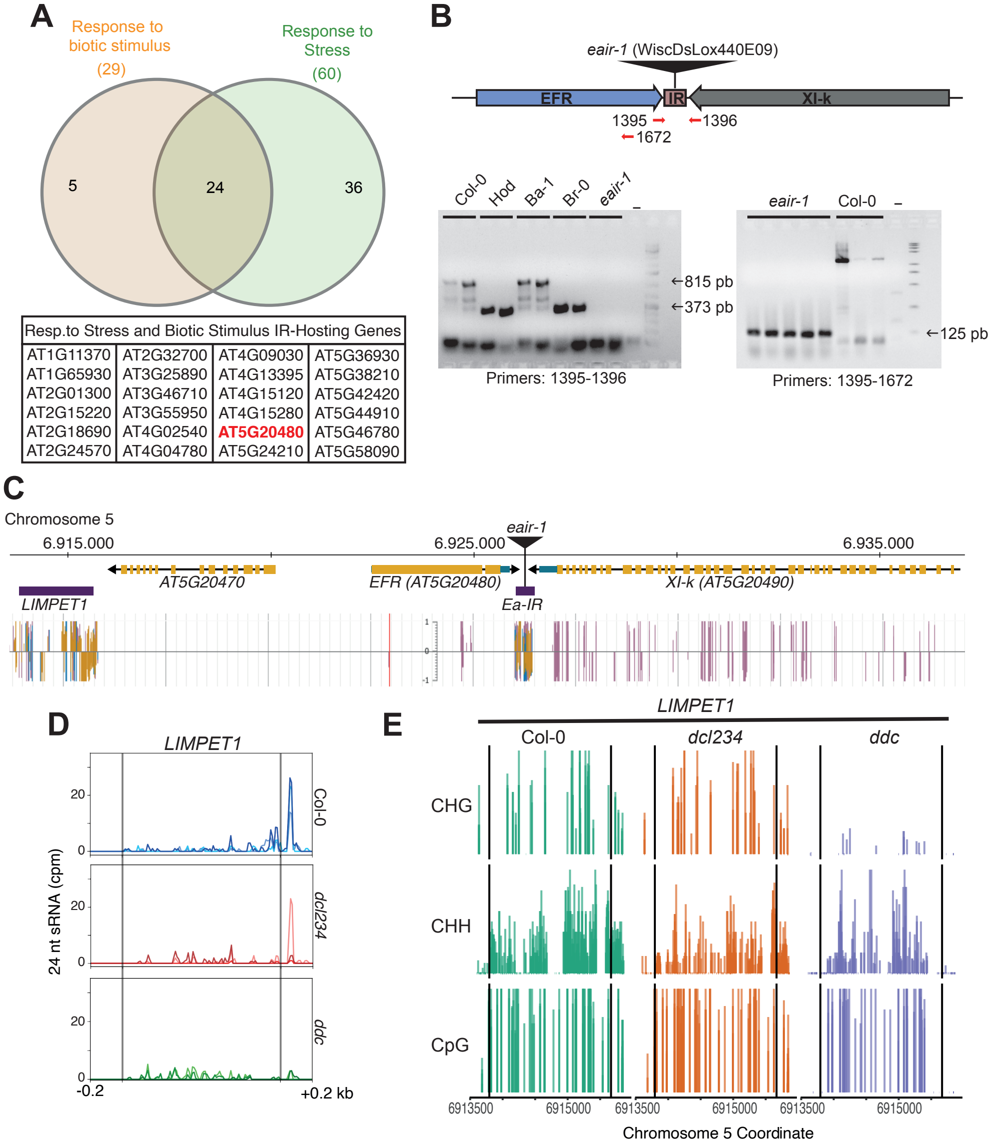


**Figure S1. Identification of an IR between *EFR* and *Xl-k*.**

(A) Gene Ontology (GO) analysis of all loci with at least one IR located 500 bp upstream or downstream of a protein-coding gene. The table at the bottom provides the gene IDs of all 24 genes in the intersection of the GO categories "response to biotic stimulus" and "response to stress". *EFR* is highlighted in red.

(B) PCR on genomic DNA to confirm the *Ea-IR* status in different *A thaliana* accession and *eair-1* T-DNA mutants. using the primers indicated above in red.

(C) Fraction of cytosine methylation in the CHH, CHG, and CG contexts across the *EFR/Xl-k* region extending to the *LIMPET1* transposon, as extracted from the Plant Epigenome Browser (https://epigenome.genetics.uga.edu/PlantEpigenome/).

(D) 24 nt siRNAs mapping to *LIMPET1* in Col-0 wild-type, *dcl234*, and *ddc* plants. Three individual sRNA-seq replicates are plotted separately and shown with different shades.

(E) Cytosine methylation in the CHG, CHH, and CG contexts at *LIMPET1* in Col-0 wild-type, *dcl234*, and *ddc* plants. Three individual BS-seq replicates are plotted for each genotype. The borders of *Ea-IR* are indicated by vertical grey lines.

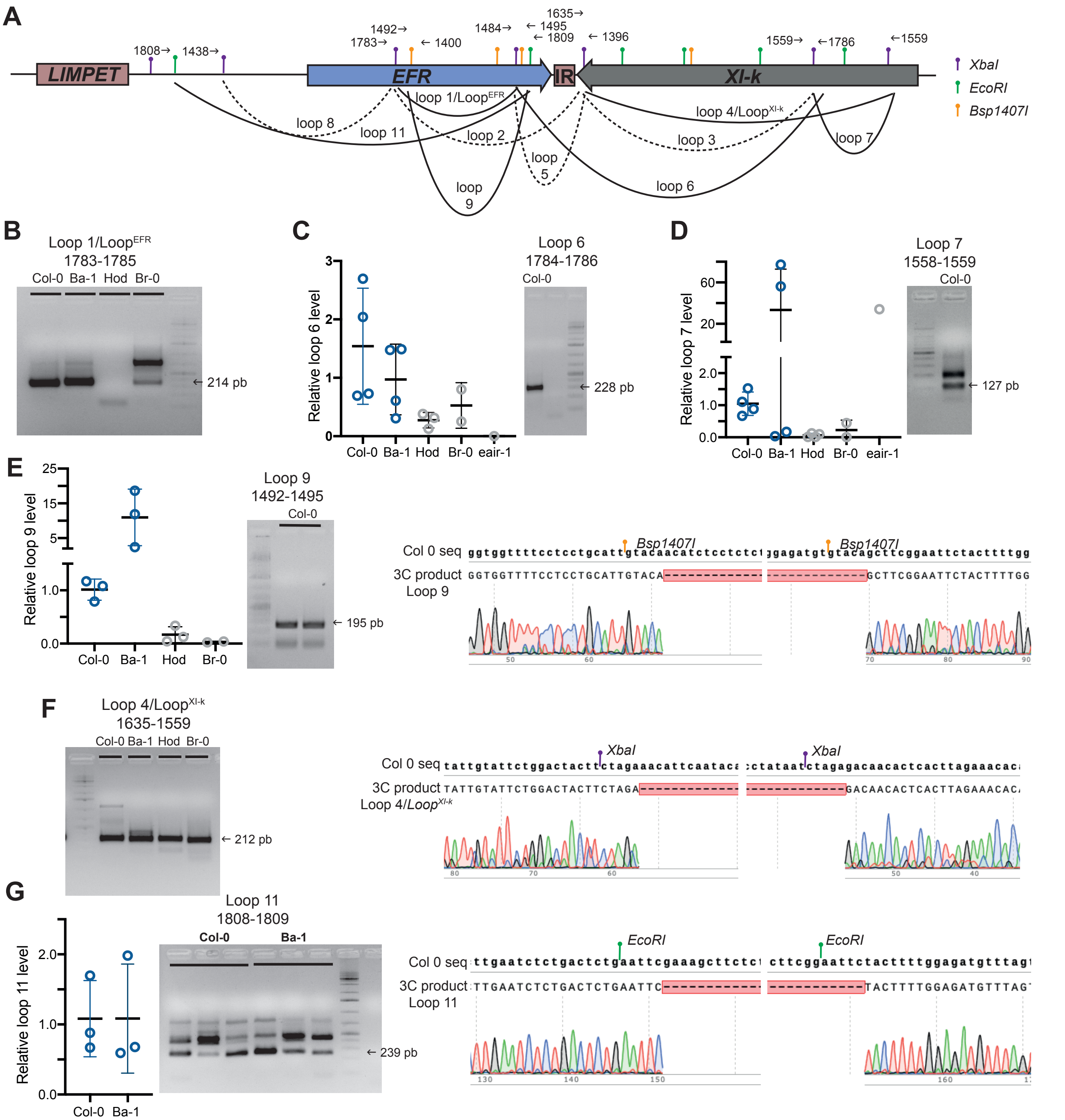


**Figure S2. Mapping of chromatin loops at the *EFR/Xl-k* locus.**
(A) Diagram of the *EFR/Xl-k* region summarizing all the interactions detected in panels B-G. The diagram includes protein coding genes and annotated transposons, the restriction sites of the enzymes used for 3C experiments (colored pins), the primers used in each experiment, detected chromatin interactions (solid curved lines), and potential chromatin interactions tested but not detected (dashed curved lines). Given the resolution of 3C experiments, the loop anchor points can be located anywhere within a ligated restriction fragment.
(B-G) Agarose gel and/or Sanger sequencing chromatograms for each loop. Primers to detect the loops are noted on top of the gel images. *Loop* 6 (C), *Loop* 7 (D), *Loop* 9 (E), and *Loop* 11 (G) abundance was measured in different accessions and mutants.

**A**

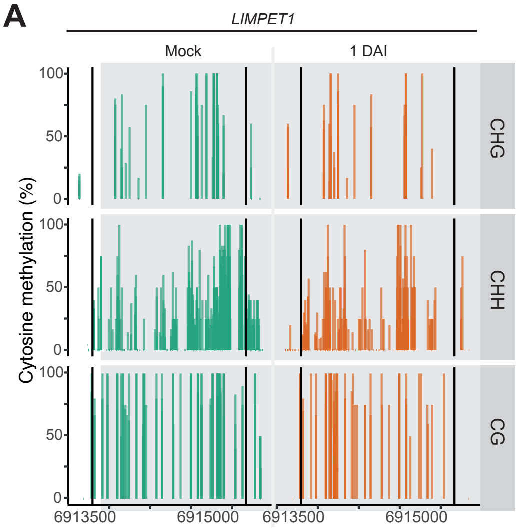


**Figure S3. Methylation at the *LIMPET1* transposon.**
Cytosine methylation in the CHG, CHH, and CG contexts at the *LIMPET1* transposon (delimited by grey vertical lines) in Col-0 wild-type plants mock-treated or infected with *Pst* DC3000 (1 dai). Three individual BS-seq replicates are plotted.

A

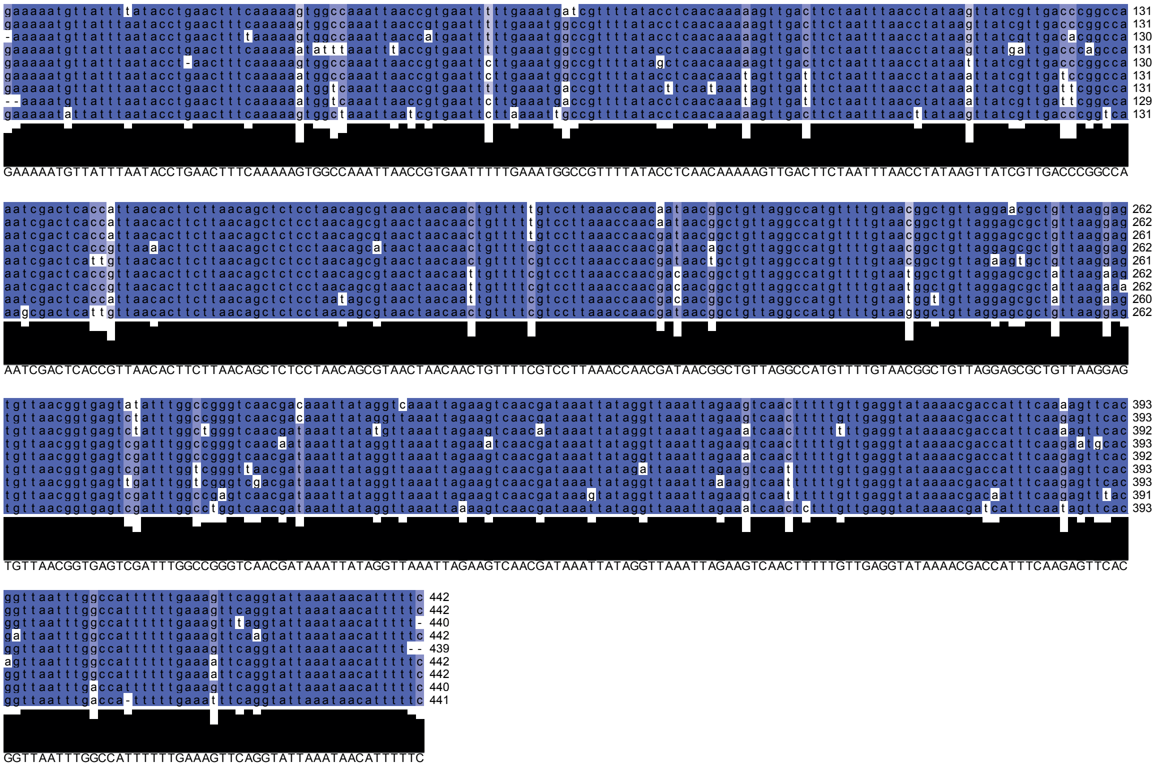


B

| Chromosome | Start | End | IR | Genotype | ID | Note | Distance | Position |
|---|---|---|---|---|---|---|---|---|
| Chr5 | 6926020 | 6926461 | Chr5_6926020_6926461 | Col_0_TAIR | AT5G20480 | EF-TU receptor | 118 | downstream |
| Chr5 | 6926020 | 6926461 | Chr5_6926020_6926461 | Col_0_TAIR | AT5G20490 | myosin family protein with Dil | 54 | downstream |
| Chr5 | 16970633 | 16971074 | Chr5_16970631_16971076 | Col_0_TAIR | AT5G42420 | Nucleotide-sugar transporter family protein | 105 | downstream |
| Chr5 | 16970633 | 16971074 | Chr5_16970631_16971076 | Col_0_TAIR | AT5G42430 | F-box and associated interaction domains-containing protein | 237 | downstream |
| Chr5 | 16970633 | 16971074 | Chr5_16970631_16971076 | Col_0_TAIR | AT5G42440 | Protein kinase superfamily protein | 2127 | downstream |
| Chr4 | 1023826 | 1024265 | Chr4_1023824_1024267 | Col_0_TAIR | AT4G02320 | Plant invertase/pectin methylesterase inhibitor superfamily | 0 | inside |
| Chr4 | 13316605 | 13317046 | Chr4_13316600_13317051 | Col_0_TAIR | AT4G26310 | elongation factor P (EF-P) family protein | 161 | upstream |
| Chr4 | 13316605 | 13317046 | Chr4_13316600_13317051 | Col_0_TAIR | AT4G26320 | arabinogalactan protein 13 | 6 | downstream |
| Chr4 | 13316605 | 13317046 | Chr4_13316600_13317051 | Col_0_TAIR | AT4G07575 | | 1199 | upstream |
| Chr4 | 13316605 | 13317046 | Chr4_13316600_13317051 | Col_0_TAIR | AT4G07585 | | 1691 | upstream |
| Chr2 | 713347 | 713788 | Chr2_713346_713789 | Col_0_TAIR | AT2G02610 | Cysteine/Histidine-rich C1 domain family protein | 527 | downstream |
| Chr1 | 4209166 | 4209607 | Chr1_4209166_4209607 | Col_0_TAIR | AT1G12360 | Sec1/munc18-like (SM) proteins superfamily | 2679 | downstream |
| Chr1 | 4209166 | 4209607 | Chr1_4209166_4209607 | Col_0_TAIR | AT1G12370 | photolyase 1 | 166 | upstream |
| Chr1 | 512695 | 513134 | Chr1_512695_513132 | Col_0_TAIR | AT1G02470 | Polyketide cyclase/dehydrase and lipid transport superfamily protein | 0 | inside |
| Chr1 | 512695 | 513134 | Chr1_512695_513132 | Col_0_TAIR | AT1G02475 | Polyketide cyclase/dehydrase and lipid transport superfamily protein | 564 | downstream |
| Chr1 | 512695 | 513134 | Chr1_512695_513132 | Col_0_TAIR | AT1G02480 | | 2359 | upstream |
| Chr1 | 20622394 | 20622834 | Chr1_20622392_20622836 | Col_0_TAIR | AT1G55270 | Galactose oxidase/kelch repeat superfamily protein | 2247 | upstream |
| Chr1 | 20622394 | 20622834 | Chr1_20622392_20622836 | Col_0_TAIR | AT1G08047 | Natural antisense transcript overlaps with AT1G55270 | 2512 | downstream |
| Chr1 | 20622394 | 20622834 | Chr1_20622392_20622836 | Col_0_TAIR | AT1G55280 | Lipase/lipooxygenase, PLAT/LH2 family protein | 95 | downstream |

C

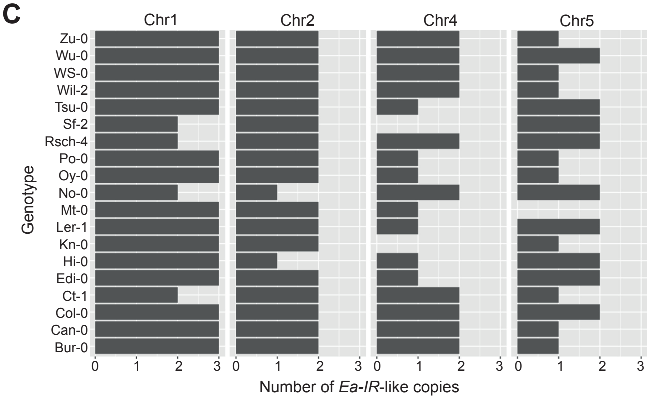


D

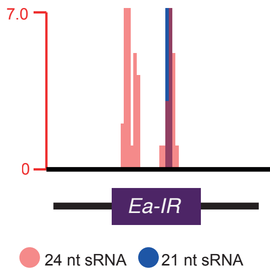

**Figure S4. *Ea-IR* related sequences in the Col-0 genome.**
(A) Nucleotide alignment of the eight *Ea-IR*-like copies in the *A. thaliana* Col-0 genome. *Ea-IR* is on top. Sequences of each *Ea-IR*-like copies are ordered as in (B) and the consensus sequence shown at the bottom.
(B) Details of each *Ea-IR*-like element indicating chromosome, start and end coordinates, complete genomic coordinates, genotype, gene ID of the closest protein coding gene, its annotation, if any, its distance to the closest *Ea-IR*-like copy in bp, and the position of the *Ea-IR*-like copy relative to the adjacent protein-coding gene.
(C) Number of *Ea-IR*-like copies on chromosomes 1, 2, 4, and 5 in 19 *A. thaliana* genome assemblies.
(D) Uniquely mapping 21 and 24 nt siRNAs over *Ea-IR* shown. Scale cpm (counts per million).

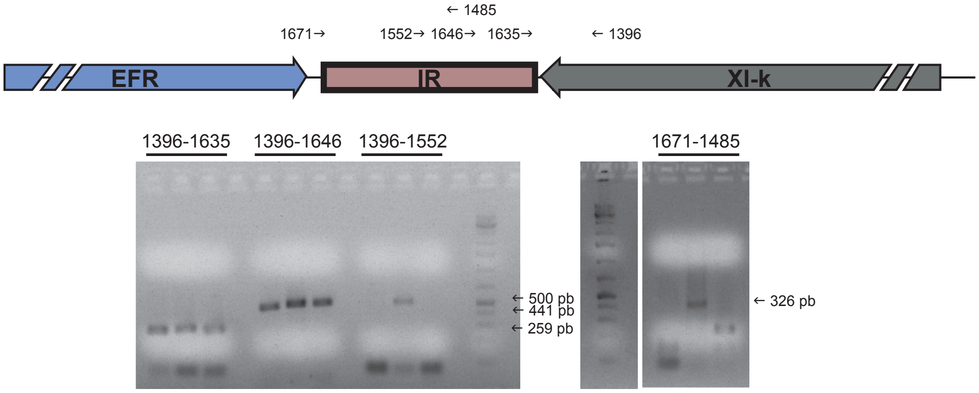


**Figure S5. Detection of long *Xl-k::Ea-IR* transcript isoform in Col-0 wild-type plants.**
Detection by RT-PCR of the long *Xl-k::Ea-IR* transcript isoform as a result of Pol II readthrough at the *Xl-k* termination site. Primers indicated on top. Results of three biological replicates are shown.

#### Supplementary Table

**Table S1.** Oligonucleotide primers used.

| Gene/element | Sequence (5'-3') | Assay | Primer Name |
|---|---|---|---|
| *ACTIN2* | GGTAACATTGTGCTCAGTGGTGG | qPCR | 189 |
| | GGAGATCCACATCTGCTGGAATG | qPCR | 190 |
| *Ea-IR* | CATTTTTCCCAATTAACATCC | qPCR/long XI-k transcript/3C | 1635 |
| | CACGTTAAAGTTCTCTGTGTCC | Genotyping | 1395 |
| | CTCATTAGAAGACAAATCAC | qPCR/long XI-k transcript | 1671 |
| | GTCCTTAAACCAACAATAAC | qPCR/long XI-k transcript | 1552 |
| | TGTTAAGGAGTGTTAACGGTG | qPCR/long XI-k transcript | 1646 |
| | GACCCGGCCAAATATACTCACCG | qPCR/long XI-k transcript | 1485 |
| | CACCCACATTAATTAACACCGAG | GFP-IR/IR-GFP cloning | 1555 |
| *EFR* | TGGCTGCAGCTAGAAGATCTGG | qPCR | 1397 |
| | TCTCTGCGGGTGTTAAAGATCTCG | qPCR | 1398 |
| | GAAAGAGTCATGTCCTGTGAC | qPCR/3C | 1808 |
| | CACCATGCAGATCTTCGTTAAG | qPCR/3C | 1783 |
| | GAGAGGTGAGATTCCAGACG | qPCR/3C | 1492 |
| | GTTGTACAATGCAGGAGGAA | qPCR/3C | 1400 |
| | GACTTTGTGTAGCTGTGGAG | qPCR/3C | 1495 |
| | CGCCTGCAAATGATTCATCTG | qPCR/3C | 1809 |
| *PR1* | ACACGTGCAATGGAGTTTGTGG | qPCR | 1631 |
| | TTGGCACATCCGAGTCTCACTG | qPCR | 1632 |
| *XI-k* | AGGGTTATGATGACAGAGGACTCG | qPCR | 1560 |
| | CTCCACCGTGAATGGAATGCTC | qPCR | 1561 |
| | GGGCGAGTTCTTATACTCGTTTTGG | qPCR/long XI-k transcript/3C/GFP-IR/IR-GFP cloning | 1396 |
| | GCTGAAGATTTAGTTTCCAG | qPCR/3C | 1559 |

| | | | |
|---|---|---|---|
| | GGATGTCTTGGAGCTCATTGAG | qPCR/3C | 1786 |
| | CCTCTAAAGAGTCTAGCAGC | qPCR/3C | 1558 |
| *Hpa ACTIN* | GTTTACTACCACGGCCGAGC | qPCR | G-44352 |
| | CGTACGGAAACGTTCATTGC | qPCR | G-44353 |
